# Prenatal Alcohol Exposure Downregulates Focal Adhesion Molecules Tensin 1 and Tensin 2 Contributing to Cerebrovascular Dysfunction

**DOI:** 10.64898/2026.10.06.757079

**Authors:** Richard Justice, Amy S. Gardiner

## Abstract

Fetal alcohol spectrum disorders (FASD) are caused by prenatal alcohol exposure (PAE), which leads to developmental defects during embryogenesis. Angiogenesis is one facet of development that is heavily dysregulated during PAE and where such impairment is poorly understood. Here we found that expression levels of focal adhesion molecules Tensin 1 (Tns1) and Tensin 2 (Tns2) were significantly reduced by alcohol exposure in brain microvascular endothelial cells (BMVECs). We also found that miR-150-5p, which we previously found was increased by PAE, regulates the expression of Tns1 and Tns2. Dual-luciferase assays demonstrated direct binding of miR-150-5p to the 3’UTRs of Tns1 and Tns2. Additionally, we utilized sprouting angiogenesis bead assays to evaluate endothelial cell sprouting in response to ethanol (EtOH) treatment and following Tns1 and/or Tns2 overexpression as well as miR-150-5p modulation. Tns1 and Tns2 rescued both EtOH- and miR-150-5p-mediated deficits in sprouting angiogenesis. Overall, we found that miR-150-5p negatively regulates Tns1 and Tns2 in BMVECs and that augmenting Tns1 and Tns2 expression may reverse the negative effects of EtOH, and subsequent miR-150-5p elevation, thus improving endothelial cell sprouting during cerebrovascular development.

## 1 Introduction

Fetal alcohol spectrum disorders (FASD) is a term used to describe various developmental disorders caused by prenatal alcohol exposure (PAE). Fetal alcohol syndrome (FAS), the most severe form of FASD, includes phenotypic changes such as a thin upper lip, smooth philtrum, and short palpebral fissure alongside alterations in neurodevelopment and behavior^1^. Many individuals with FASD lack recognizable physical features and often go undiagnosed, highlighting the need to understand the mechanisms underlying FASD-associated neurodevelopmental deficits^2,3^.

Cerebrovascular dysfunction is one potential contributor to FASD-associated neurodevelopmental deficits, particularly during development of the fetal brain^4,5^. Sprouting angiogenesis is a fundamental mechanism of brain vascularization during development, characterized by the migration of endothelial tip cells at the leading edge of nascent vessel sprouts, followed by proliferating stalk cells that contribute to lumen formation^6,7^. Subsequently, neighboring sprouts undergo anastomosis to generate a functional and interconnected vascular network^6,8,9^. In addition to the reported cerebrovascular alterations following PAE^4^ acute alcohol exposure has been shown to be associated with impaired angiogenesis in wound healing models, alongside extracellular matrix (ECM) collagen content, and lower fibroblast growth factor (FGF) and vascular endothelial growth factor (VEGF) expression^10^. During angiogenesis, focal adhesions enable endothelial cells to generate the traction forces needed to migrate through the ECM and perform mechanotransduction^9,11–13^. Tensin 1 (Tns1) and Tensin 2 (Tns2) are focal adhesion proteins involved in cell proliferation, adhesion, and signaling^14,15^. Tns1 localizes to both focal and fibrillar adhesions, whereas Tns2 is found predominantly in focal adhesions^16^. Both proteins contain several functional domains such as Phosphatase and Tensin Homologue (PTEN), C2, Actin Binding domain (ABD), Src Homology 2 (SH2), and Phosphotyrosine Binding Domain (PTB)^15^. These domains allow tensins to interact with actin and integrins, supporting cell-ECM signaling and adhesion^17^. In particular the SH2 domain binds phosphotyrosine-containing proteins and have been implicated in PI3K/Akt and β-integrin/FAK signaling pathways^18^.

Little is known about the mechanisms that contribute to the pathogenesis of PAE-mediated neurovascular dysfunction; however, these alterations may be explained by post-transcriptional gene regulation through microRNAs (miRNAs). miRNAs are small non-coding single-stranded RNA molecules 18-22 nucleotides in length that bind to the 3’ untranslated region (3’UTR) of a target messenger RNA (mRNA) and induce mRNA degradation or inhibit translation^19–21^. One miRNA of interest is miR-150-5p, which has been shown to negatively regulate angiogenesis in the brain^22,23^. Previous work from our lab showed that miR-150-5p is upregulated in primary brain microvascular endothelial cells (BMVECs) isolated from embryonic day 18 (E18) PAE cortices compared to saccharin (SAC)-exposed controls^24^. Recently, we found that TET-mediated DNA demethylation was enhanced at the miR-150-5p promoter, increasing its promoter activity in alcohol-exposed BMVECs^25^. Additionally, we demonstrated that miR-150-5p inhibits vascular endothelial zinc finger 1 (Vezf1), reducing the migration and tube formation of BMVECs^24^.

In this study, we show that expression levels of Tns1 and Tns2 are reduced by alcohol exposure in BMVECs, and that both molecules are regulatory targets of miR-150-5p. Using a sprouting angiogenesis bead assay, we modeled the anti-angiogenic effects of both alcohol exposure and miR-150-5p on BMVECs. Moreover, overexpression of Tns1 and Tns2 restored the angiogenic function of BMVECs after alcohol exposure and miR-150-5p overexpression. Our findings provide mechanistic insight into how alcohol and miRNAs contribute to vascular dysfunction, which may inform the development of future FASD therapeutics.

## 2 Materials and Methods

### 2.1 PAE Mouse Model

A previously described moderate drinking paradigm was used^24^. Female C57BL/6J mice were given access to 10% (w/v) ethanol (EtOH) in 0.066% (w/v) saccharin (SAC) or 0.066% SAC-sweetened water for 4hrs daily. Dams had access to these solutions 4hrs every day before breeding to establish a consistent drinking pattern. Embryos were harvested at E18 for further studies.

### 2.2 Cell culture and transfections

Immortalized murine BMVECs (bEnd.3) and HeLa cells were purchased from Sigma-Aldrich and were grown in Dulbecco’s Modified Eagle Medium (DMEM, Corning #14224009) supplemented with 10% fetal bovine serum (Neuromics, #N23H23), 200 mM L-glutamine (Gibco, #2660218) and 1% penicillin-Streptomycin (Gibco, #15140-122). Cells were incubated at 37°C and 5% CO_2_. Overexpression or inhibition of miR-150-5p was performed by transfecting bEnd.3 BMVECs or HeLa cells with 25nM of miRCURY LNA™ miR-150-5p mimics, miR-150-5p inhibitors, negative control mimics, or negative control inhibitors (Qiagen). Overexpression of Tns1 (Addgene, plasmid #105297) or Tns2 (Addgene, plasmid #105298) was achieved by transfecting 1µg (12-well plate) or 5µg (6-well plate). The bEnd.3 Cell Avalanche® Transfection Reagent (EZ Biosystems) or Lipofectamine™ 2000 (Thermo Fisher Scientific) was used according to manufacturer instructions.

### 2.3 Cell model of moderate alcohol exposure

We utilized a previously established cell model of alcohol exposure^24,25^. Briefly, bEnd.3 BMVECs were treated with a 25mM EtOH-media solution for 24hrs, and media was replaced every 12hrs. To prevent EtOH evaporation, a reservoir of 50 mM EtOH was placed in the 37°C, 5% CO_2_ incubator. Following EtOH treatment, cells were used for RT-qPCR, or in preparation for bead assays.

### 2.4 RNA extraction and RT-qPCR

Total RNA was extracted from treated bEnd.3 cells using Trizol. RNA was quantified using the Qubit™ 4 Fluorometer (Thermo Fisher Scientific). cDNA synthesis was performed using the SuperScript™ IV First-Strand Synthesis System (Thermo Fisher Scientific) with Oligo(dT) primers. Quantitative PCR (qPCR) was performed using the Power SYBR™ Green PCR Master Mix (Applied Biosystems) with primers for mouse GAPDH, Tns1, or Tns2; additionally, human Tns1 and Tns2 primers were used to confirm plasmid overexpression. Real-time PCR was performed on the AriaMx Real-Time PCR System (Agilent). Relative quantification of gene expression was performed using 2^(−ΔΔCt)^ calculations where mRNA expression levels were normalized to GAPDH.

### 2.5 Dual-luciferase assays

A ∼2900bp (Tns1) and ∼300bp (Tns2) segment of the 3’UTR containing the miR-150-5p binding sight was cloned into the pISO firefly luciferase reporter (Addgene, plasmid #12178)) using *SacI* and *XbaI* restriction sites from gDNA extracted from BMVECs. The DNA fragment was PCR amplified using the following primers: Tns1 Forward 5’-GAAGAGCTCAGTGCAATGTTGGGAGAGGA-3’, Reverse 5’-GGGTCTAGAACTGTTAGTTCTCCCAGCCT-3’; Tns2 Forward 5’-GAAGAGCTCACAAGCTCCGAGTGCACA-3’ Reverse 5’-GAGTCTAGAGTGGAGGGAAGGCAGGAAG-3’. HeLa cells were transfected with the Tns1 or Tns2 3’UTR construct along with the pIS2 Renilla control vector (Addgene, plasmid #12177). The cells were also co-transfected with 25nM miR-150-5p miRCURY LNA™ miR-150-5p mimics or control mimics (Qiagen). Dual-luciferase assays were performed using the Dual-Luciferase® Reporter Assay System (Promega Corporation) and luciferase activity was measured using the Infinite® 200 plate reader (Tecan).

### 2.6 Sprouting angiogenesis bead assays

BMVECs were incubated at 400 cells/Cytodex™ 3 microcarrier bead (Cytiva) for 4hr and agitated every 20min in endothelial cell media (ScienCell). Cell-coated beads (CCB) were transferred to a 6-well plate to isolate CCB from free cells. Cell-coated beads were later imbedded in a fibrin clot supplemented with of 2 mg/mL Fibrinogen (Sigma-Aldrich), 0.15 U/mL aprotinin (Sigma-Aldrich), 5 ng/mL VEGF (ThermoFisherScientific), and 5 ng/mL FGF (ThermoFisherScientific). The fibrin clot was formed by mixing 500µL of the fibrinogen mix with 12.5µL thrombin at 50 U/mL (Sigma-Aldrich) in a 24-well plate then incubated at 37°C for 15min. Endothelial cell media was added to each well and sprouts were imaged using the Echo Revolve (Discover Echo) after 24hr. Sprouting was quantified using WimSprout (Wimasis). The sprouting angiogenesis bead assay was adapted from Clavane *et.al*.^26^

### 2.7 Statistical analysis

Statistical analyses for all data were performed with at least three independent replicates using GraphPad Prism v11.0 software (GraphPad Software Inc., USA).

## 3 Results

### 3.1 Tensin 1 and Tensin 2 expression levels are reduced by alcohol exposure in brain microvascular endothelial cells

Previously we performed digital spatial profiling of E18 PAE and SAC cortices following moderate PAE^27^. Using this data, we found that Tns1 and Tns2 levels were significantly reduced in the E18 PAE BMVECs compared to SAC controls (Figure 1A, B). We replicated these results in our cell model of moderate alcohol exposure, where bEnd.3 BMVECs are treated with 25mM EtOH for 24hrs^24^. Using RT-qPCR, we found that Tns1 and Tns2 expression levels were significantly downregulated in EtOH-treated BMVECs compared to untreated controls (Figure 1C, D).

**Figure 1:**
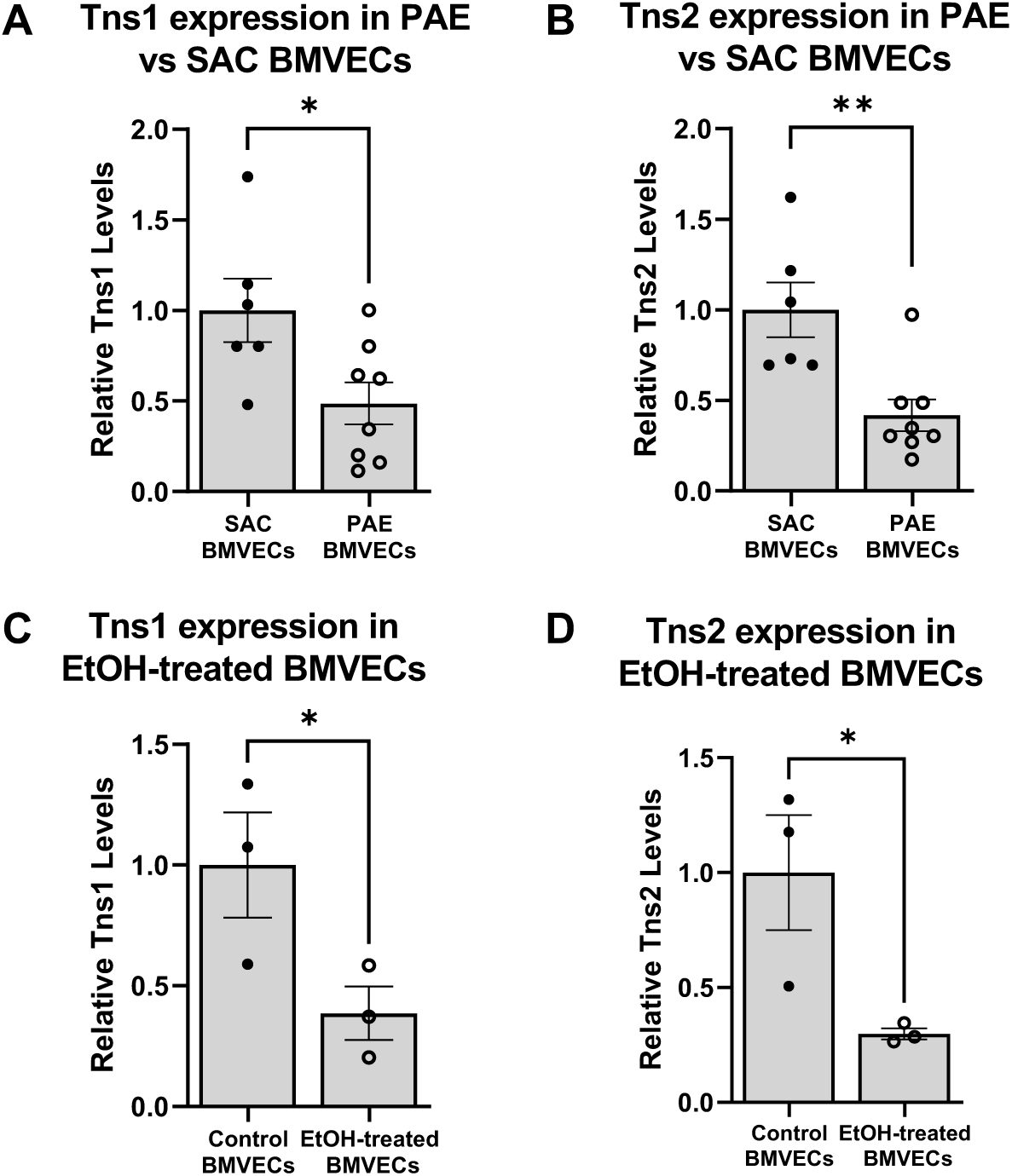
Tensin 1 and Tensin 2 expression levels are reduced by alcohol exposure in brain microvascular endothelial cells. **(A)** Tns1 expression from DSP data of PAE vs SAC BMVECs. n=6-8 ROIs. p=0.0254. **(B)** Tns2 expression from DSP data of PAE vs SAC BMVECs. n=6-8 ROIs. p=0.0042. **(C)** Tns1 expression in EtOH-treated vs untreated BMVECs. RT-qPCR. n=3. p=0.0331. **(D)** Tns2 expression in EtOH-treated vs untreated BMVECs. RT-qPCR. n=3. p=0.0248. Error bars=SEM. *p<0.05, **p<0.01.

### 3.2 Tensin 1 and Tensin 2 are molecular targets of miR-150-5p

Using TargetScan Mouse Release 8.0, we discovered that Tns1 and Tns2 are putative regulatory targets of miR-150-5p, containing conserved miR-150-5p-binding sites in their 3’UTRs (Figure 2A). To evaluate the effect of miR-150-5p on Tns1 and Tns2, we transfected BMVECs with miR-150-5p LNA™ mimics or inhibitors, and their respective controls, and measured gene expression using RT-qPCR. Overexpression of miR-150-5p decreased both Tns1 and Tns2 levels compared to the negative mimic control (Figure 2B,D). In addition, the knockdown of miR-150-5p significantly increased both Tns1 and Tns2 expression (Figure 2C,E). Overexpression and inhibition of miR-150-5p was confirmed using RT-qPCR (Supplementary Figure 1A,B).

**Figure 2:**
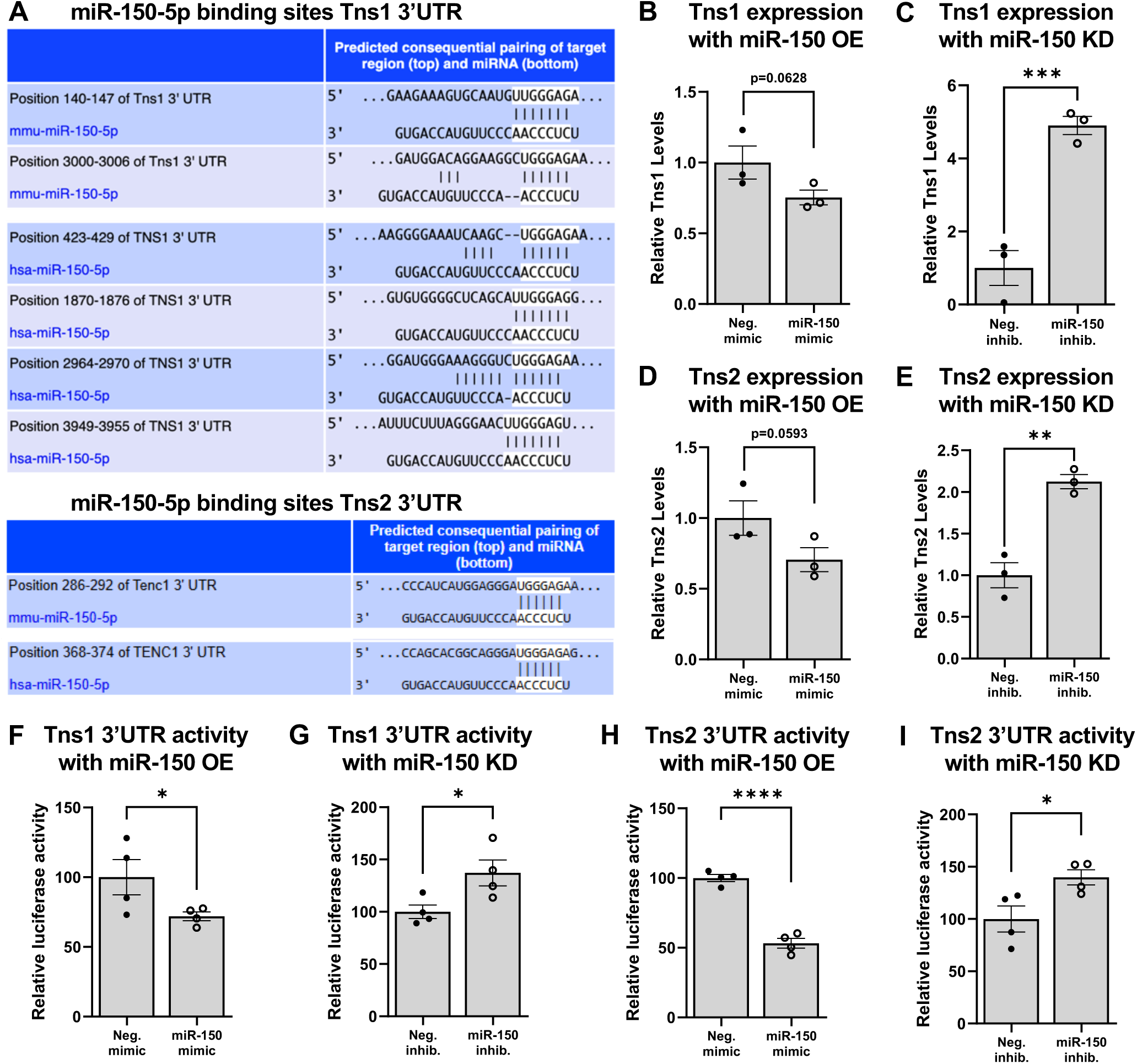
miR-150-5p binds the Tensin 1 and Tensin 2 3’UTRs and regulates their expression. **(A)** Tns1 (top) and Tns2/Tenc1 (bottom) 3’UTRs contain conserved miR-150-5p-binding sites between mouse (mmu-miR-150-5p) and human (hsa-miR-150-5p). **(B,C)** Tns1 expression following miR-150-5p OE (p=0.0628) and KD (p=0.0010) in BMVECs. RT-qPCR. n=3. **(D,E)** Tns2 expression following miR-150-5p OE (p=0.0594) and KD (p=0.0014) in BMVECs. RT-qPCR. n=3. **(F,G)** Luciferase activity of a Tns1 3’UTR firefly reporter following miR-150-5p OE (p= 0.0376) and KD (p=0.0189). n=4. **(H,I)** Luciferase activity of a Tns2 3’UTR firefly reporter following miR-150-5p OE (p<0.0001) and KD (p=0.0160). n=4. Error bars=SEM. *p<0.05, ****p<0.0001.

We also evaluated the direct binding of miR-150-5p to the 3’UTRs of Tns1 and Tns2 using a dual luciferase assay. We cloned ∼2900 segment of Tns1 and ∼300 segment of Tns2 containing the miR-150-5p binding site into the pISO firefly luciferase reporter. The Tns1 and Tns2 constructs were co-transfected with the miR-150-5p LNA™ mimics or inhibitors. Compared to their negative controls, the miR-150-5p mimics significantly decreased luciferase activity for both Tns1 and Tns2 (Figure 2F,H). Conversely, inhibition of miR-150-5p with Tns1 or Tns2 constructs significantly increased luciferase activity compared to their respective controls (Figure 2G,I).

### 3.3 Tensin 1 and Tensin 2 overexpression rescues EtOH-mediated deficits in angiogenic sprouting

To evaluate the effect of alcohol exposure on endothelial cell sprouting, we used a sprouting angiogenesis bead assay^26^. bEnd.3 cells were transfected with expression vectors for the following: Tns1, Tns2, Tns1 and Tns2 in combination (Tns1/2), or pcDNA3 as a control. Following EtOH treatment, BMVECs were allowed to adhere to the beads and treated with angiogenic factors to promote sprouting. After 24 hrs, beads were imaged using the Echo Revolve microscope (Figure 3A). Images were submitted to Wimasis for analysis via the WimSprout program and vascular parameters were quantified (Figure 3B), including the number of sprouts, sprout area (px), mean sprout length (px), and cumulative sprout length (px). We found that EtOH treatment significantly decreased the number of sprouts, sprout area, mean sprout length, and cumulative sprout length compared to untreated controls (Figure 3C-F). Additionally, EtOH-treated BMVECs transfected with the Tns1-, Tns2-, and Tns1/2-expressing plasmids significantly increased all vascular parameters compared to the EtOH-treated pcDNA3 control (Figure 3C-F). Altogether, these data showed that Tns1 and Tns2, alone or in combination, rescued the negative effects of EtOH treatment on BMVEC angiogenic sprouting.

**Figure 3:**
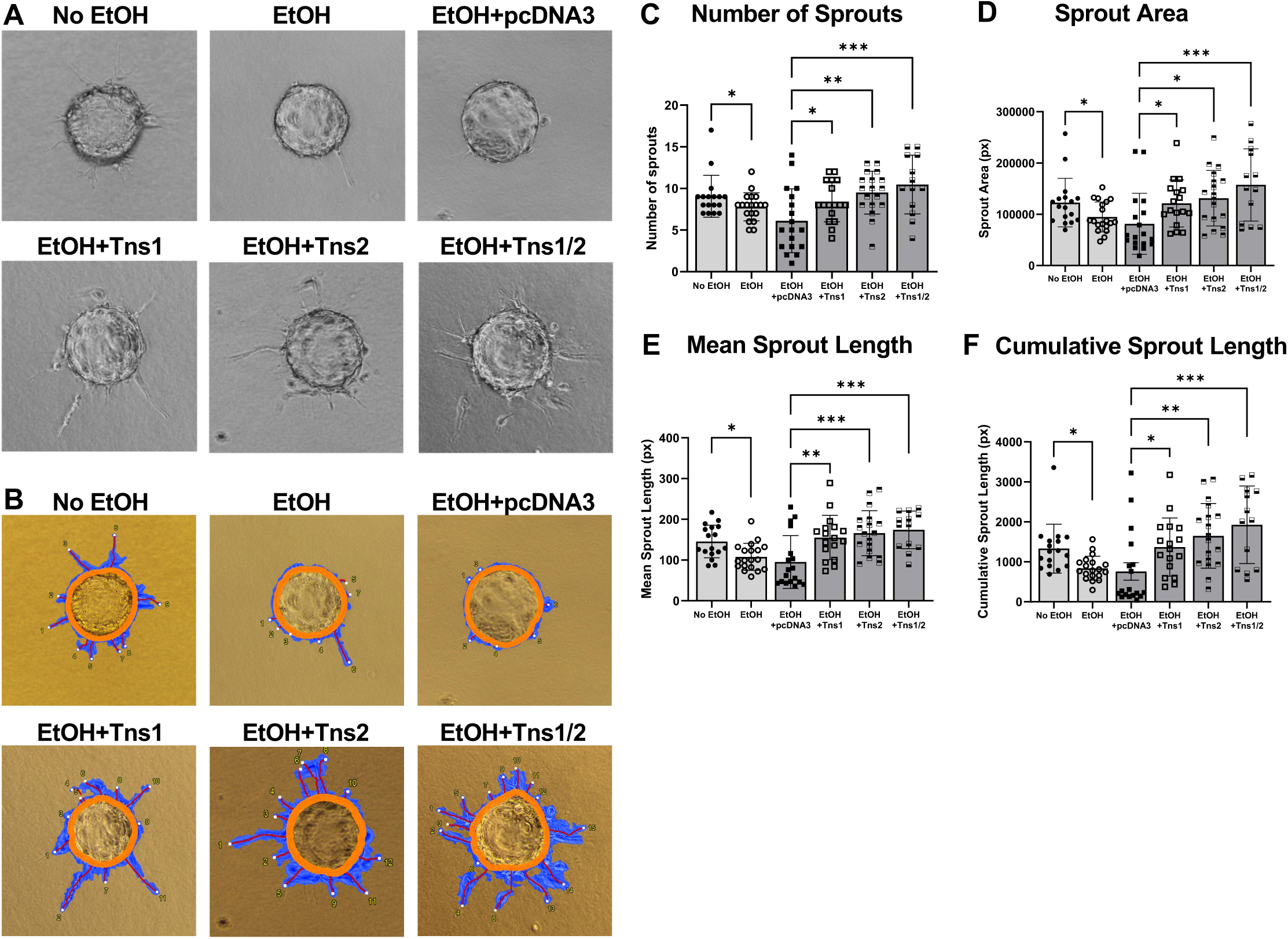
Tensin 1 and Tensin 2 rescues sprouting angiogenesis following alcohol exposure. **(A)** Representative images of cell-coated beads from sprouting angiogenesis bead assays where BMVECs were treated with EtOH and transfected with Tns1 and/or Tns2 expressing plasmids. **(B)** Corresponding images to (A) of cell-coated beads from sprouting angiogenesis bead assays with the WimSprout mask (Wimasis). **(C)** Quantification of number of sprouts: EtOH vs No EtOH (p=0.0406); EtOH+Tns1 vs EtOH+pcDNA3 (p=0.0333); EtOH+Tns2 vs EtOH+pcDNA3 (p=0.0019); EtOH+Tns1/2 vs EtOH+pcDNA3 (p=0.0003). **(D)** Quantification of sprout area: EtOH vs No EtOH (p=0.0188); EtOH+Tns1 vs EtOH+pcDNA3 (p=0.0438); EtOH+Tns2 vs EtOH+pcDNA3 (p=0.0114); EtOH+Tns1/2 vs EtOH+pcDNA3 (p=0.0006). **(E)** Quantification of mean sprout length: EtOH vs No EtOH (p=0.0021); EtOH+Tns1 vs EtOH+pcDNA3 (p=0.0024); EtOH+Tns2 vs EtOH+pcDNA3 (p=0.0003); EtOH+Tns1/2 vs EtOH+pcDNA3 (p=0.0003). **(F)** Quantification of cumulative sprout length: EtOH vs No EtOH (p=0.0019); EtOH+Tns1 vs EtOH+pcDNA3 (p=0.0398); EtOH+Tns2 vs EtOH+pcDNA3 (p=0.0028); EtOH+Tns1/2 vs EtOH+pcDNA3 (p=0.0004). n=13-19 images (beads)/condition. Error bars=SD. *p<0.05, **p<0.01, ***p<0.001.

### 3.4 Tensin 1 and Tensin 2 overexpression reverses miR-150-5p-mediated defects in angiogenic sprouting

To understand how overexpression of Tns1 and Tns2 affect miR-150-5p-associated vascular defects, we first transfected miR-150-5p mimics or inhibitors into bEnd.3 BMVECs and performed the sprouting angiogenesis bead assay (Figure 4A,B). Compared to their controls, miR-150-5p overexpression significantly decreased the number of sprouts, sprout area, mean sprout length, and cumulative sprout length in BMVECs, and miR-150-5p inhibition significantly increased these parameters (Figure 4C-F). We then examined the effect of miR-150-5p inhibition in combination with EtOH treatment (Figure 5A,B). Compared to the respective control, EtOH-treated cells transfected with miR-150-5p inhibitors significantly increased the number of sprouts, sprout area, and cumulative sprout length (Figure 5C-F). Finally, we assessed whether Tns1 or Tns2 could counter the effects of miR-150-5p overexpression of BMVEC sprouting (Figure 6A,B). miR-150-5p-overexpressing BMVECs transfected with the Tns1-, Tns2-, and Tns1/2-expressing plasmids significantly increased the number of sprouts, sprout area, mean sprout length, and cumulative sprout length compared to the miR-150-5p-overexpressing BMVECs transfected with pcDNA3 control (Figure 6C-F). Altogether, these data showed that Tns1 and Tns2, alone or in combination, reversed the negative effects of EtOH treatment on BMVEC angiogenic sprouting.

**Figure 4:**
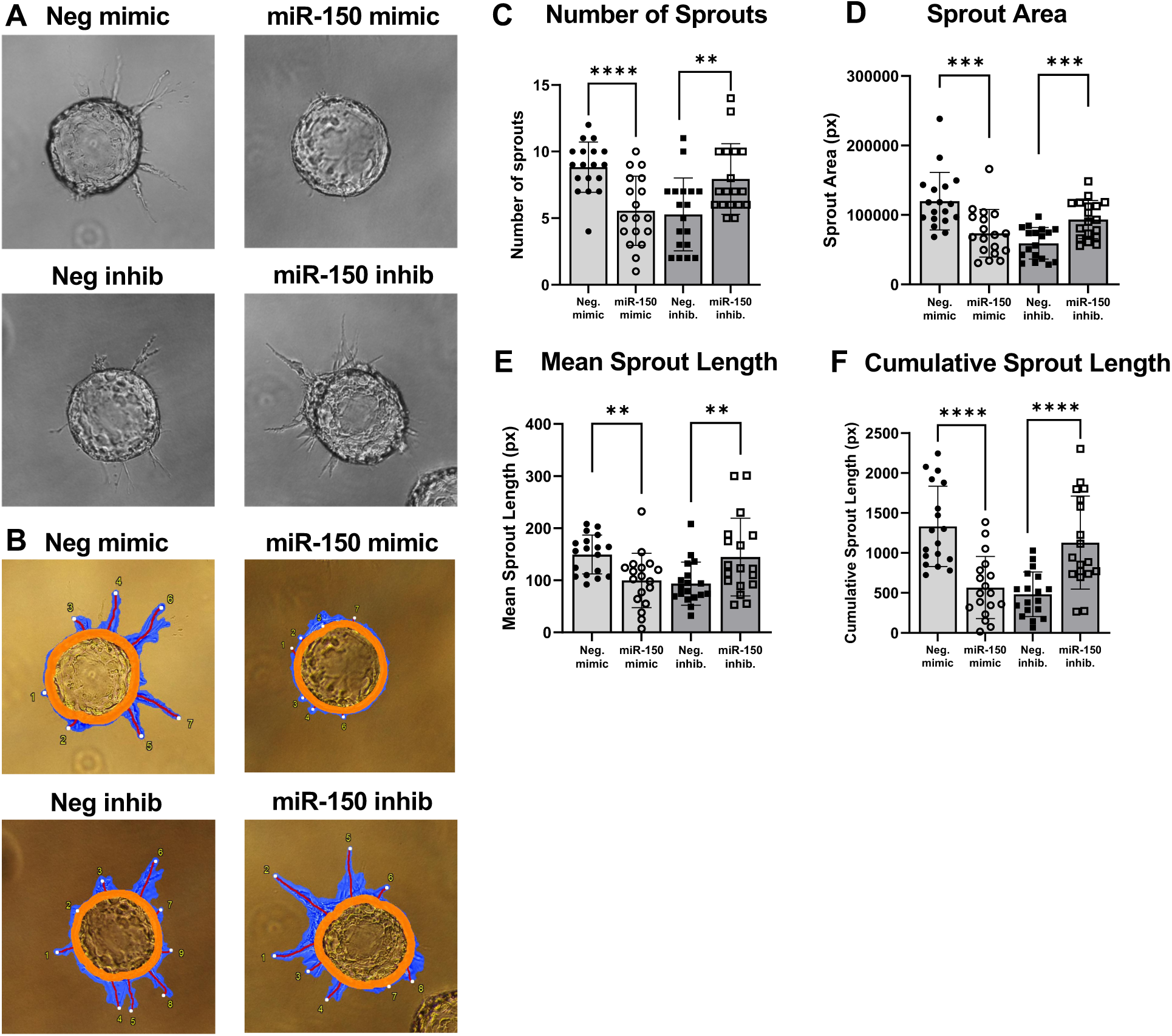
miR-150-5p reduces sprouting angiogenesis. **(A)** Representative images of cell-coated beads from sprouting angiogenesis bead assays where BMVECs were transfected with miR-150-5p mimics, inhibitors, or their respective controls. **(B)** Corresponding images to (A) of cell-coated beads from sprouting angiogenesis bead assays with the WimSprout mask. **(C)** Quantification of number of sprouts: miR-150-5p mimic vs negative control mimic (p<0.0001); miR-150-5p inhibitor vs negative control inhibitor (p=0.0027). **(D)** Quantification of sprout area: miR-150-5p mimic vs negative control mimic (p=0.0004); miR-150-5p inhibitor vs negative control inhibitor (p=0.0001). **(E)** Quantification of mean sprout length: miR-150-5p mimic vs negative control mimic (p=0.0011); miR-150-5p inhibitor vs negative control inhibitor (p=0.0079). **(F)** Quantification of cumulative sprout length: miR-150-5p mimic vs negative control mimic (p<0.0001); miR-150-5p inhibitor vs negative control inhibitor (p<0.0001). n=18 images (beads)/condition. Error bars=SD. *p<0.05, **p<0.01, ***p<0.001, ****p<0.0001.

**Figure 5:**
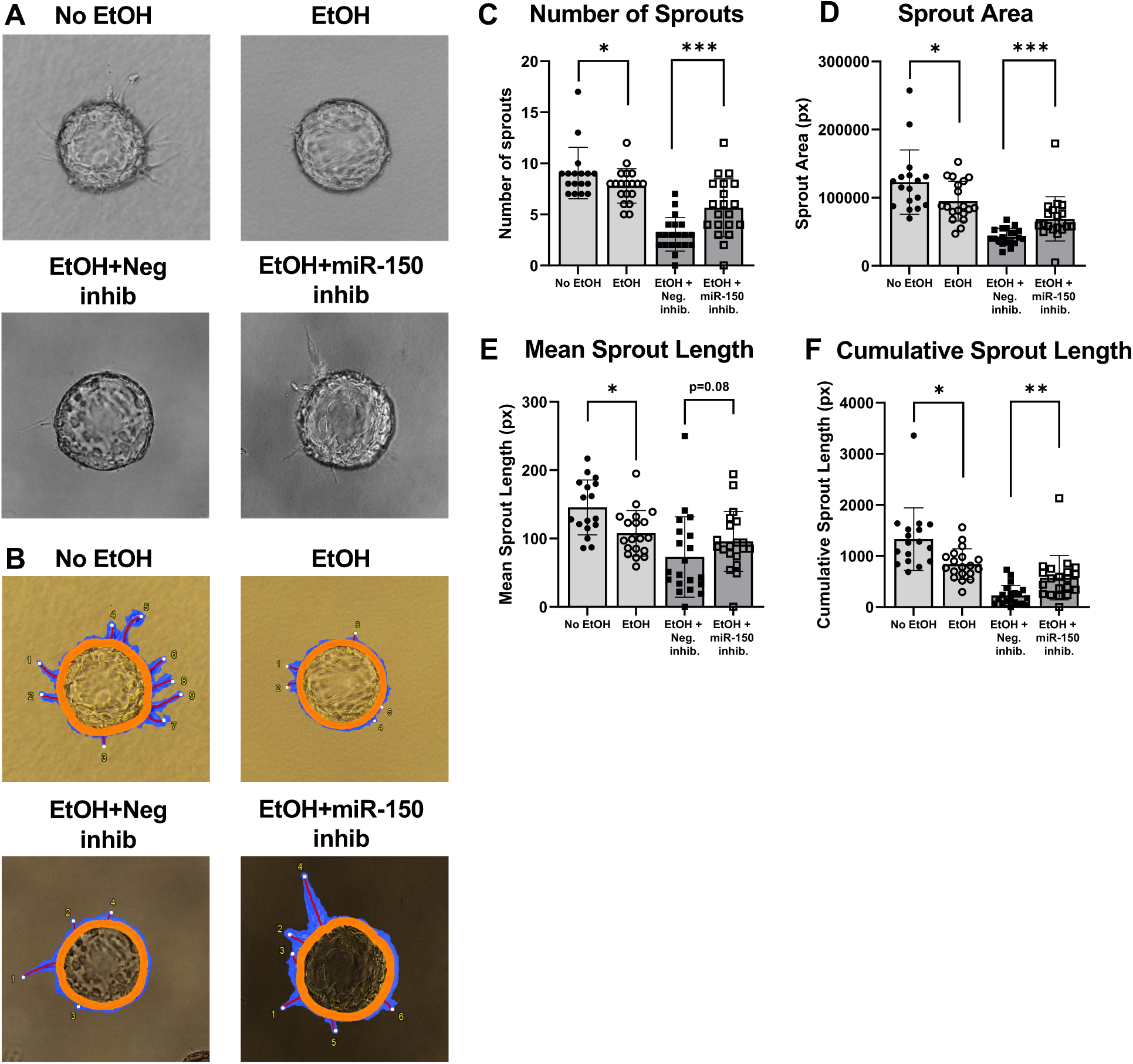
miR-150-5p inhibition restores sprouting angiogenesis following alcohol exposure. **(A)** Representative images of sprouting angiogenesis bead assays where bEnd.3 cells were treated with EtOH and transfected with miR-150-5p inhibitors or the respective control. **(B)** Corresponding images to (A) of cell-coated beads from sprouting angiogenesis bead assays with the WimSprout mask. **(C)** Quantification of number of sprouts: EtOH vs No EtOH (p=0.0406); EtOH+miR-150-5p inhibitor vs EtOH+negative control inhibitor (p=0.0005). **(D)** Quantification of sprout area: EtOH vs No EtOH (p=0.0188); EtOH+miR-150-5p inhibitor vs EtOH+negative control inhibitor (p=0.0009). **(E)** Quantification of mean sprout length: EtOH vs No EtOH (p=0.0021); EtOH+miR-150-5p inhibitor vs EtOH+negative control inhibitor (p=0.0867). **(F)** Quantification of cumulative sprout length: EtOH vs No EtOH (p=0.0019); EtOH+miR-150-5p inhibitor vs EtOH+negative control inhibitor (p=0.0013). n=17-20 images (beads)/condition. Error bars=SD. *p<0.05, **p<0.01, ***p<0.001.

**Figure 6:**
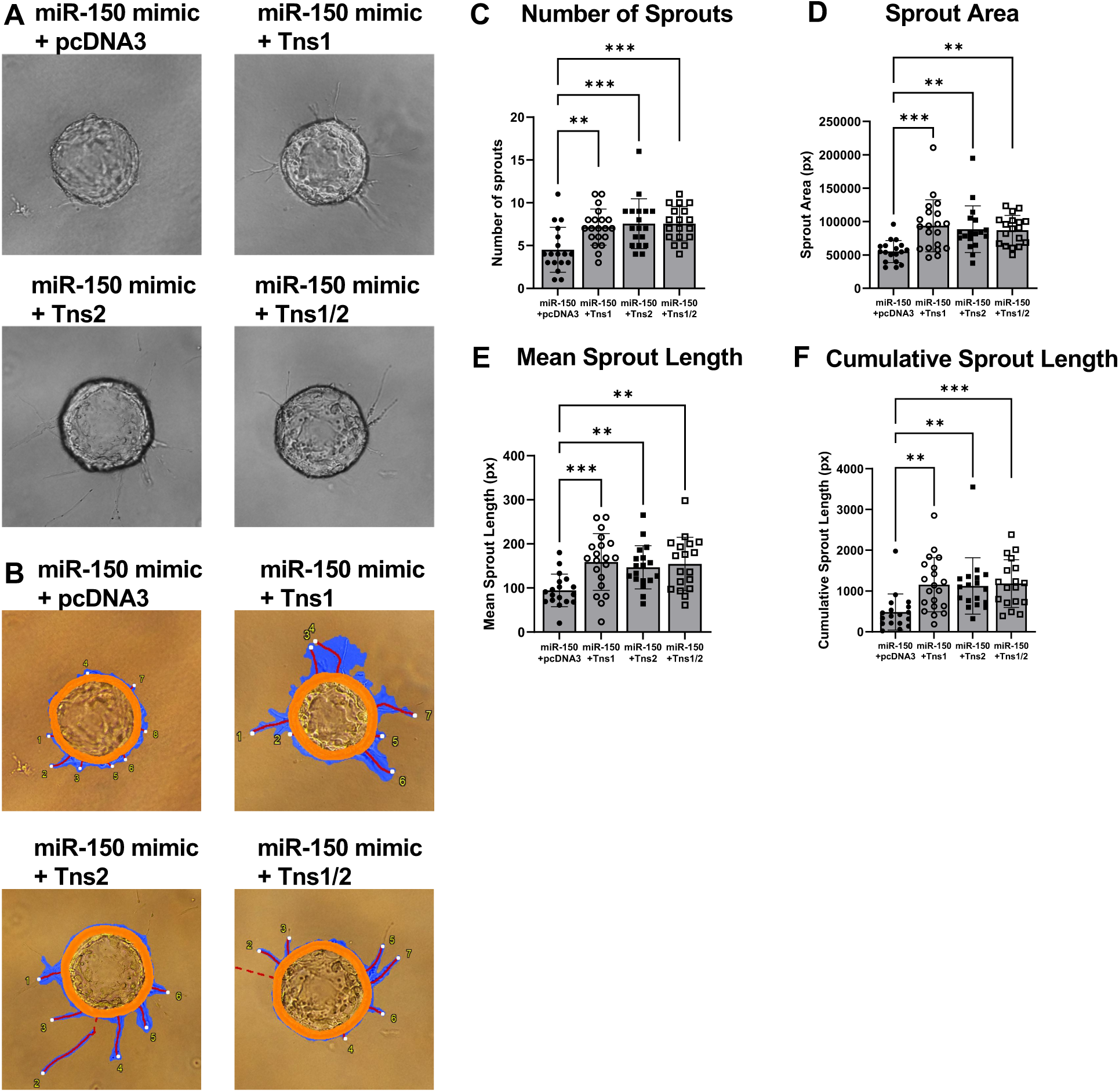
Tensin 1 and Tensin 2 reverses the miR-150-5p-mediated reduction of sprouting angiogenesis. **(A)** Representative images of sprouting angiogenesis bead assays where bEnd.3 cells were co-transfected with Tns1 and/or Tns2 expressing plasmids and miR-150-5p mimics or the respective control. **(B)** Corresponding images to (A) of cell-coated beads from sprouting angiogenesis bead assays with the WimSprout mask. **(C)** Quantification of number of sprouts: miR-150-5p+Tns1 vs miR-150-5p+pcDNA3 (p=0.0013); miR-150-5p+Tns2 vs miR-150-5p+pcDNA3 (p=0.0003); miR-150-5p+Tns1/2 vs miR-150-5p+pcDNA3 (p=0.0003). **(D)** Quantification of sprout area: miR-150-5p+Tns1 vs miR-150-5p+pcDNA3 (p=0.0002); miR-150-5p+Tns2 vs miR-150-5p+pcDNA3 (p=0.0013); miR-150-5p+Tns1/2 vs miR-150-5p+pcDNA3 (p=0.0020). **(E)** Quantification of mean sprout length: miR-150-5p+Tns1 vs miR-150-5p+pcDNA3 (p=0.0005); miR-150-5p+Tns2 vs miR-150-5p+pcDNA3 (p=0.0048); miR-150-5p+Tns1/2 vs miR-150-5p+pcDNA3 (p=0.0014). **(F)** Quantification of cumulative sprout length: miR-150-5p+Tns1 vs miR-150-5p+pcDNA3 (p=0.0011); miR-150-5p+Tns2 vs miR-150-5p+pcDNA3 (p=0.0022); miR-150-5p+Tns1/2 vs miR-150-5p+pcDNA3 (p=0.0009). n=18-20 images (beads)/condition. Error bars=SD. *p<0.05, **p<0.01, ***p<0.001.

## 4 Discussion

The prevalence of FASD in the U.S. is high, with an estimated 1-5% of first grade students affected by some form of FASD^28^. Children born with FASD are subject to lifelong neurodevelopmental and behavioral effects, often apparent in adolescence where individuals display higher incidence of psychopathology, attention defects, and impulsiveness^29,30^; with additional traits of larger total cerebral volume and regional cortical volume relative to unexposed individuals^31^. PAE contributes to the pathophysiology of the developing cerebral vascular network, demonstrated in Jégou *et al*., where prenatal alcohol exposed P2 pups displayed a significantly disorganized vasculature compared to controls, along with a reduction in VEGF-R2 protein levels^4^.

The molecular underpinnings of PAE-mediated cerebrovascular alterations is an area of study that needs more attention. Our approach of utilizing miRNA inhibitors to attenuate cerebrovascular defects in PAE is a promising pathway for future studies, as we have shown that miR-150-5p is a negative regulator of angiogenesis^24,32^. We further characterized the PAE-mediated regulation of miR-150-5p, showing that miR-150-5p is upregulated in part by demethylation at its promoter, through the activity of the Tet1 demethylase^25^.

Our current study investigated the PAE-mediated regulation of Tns1 and Tns2 expression and how their downregulation contributes to a reduction in BMVEC angiogenic sprouting. We found that their downregulation is facilitated by miR-150-5p, as both Tns1 and Tns2 are molecular targets of miR-150-5p through the direct binding of the 3’UTRs of both transcripts. Previous studies found Tns1 to be a target of other microRNAs such as miR-522-3p^33^, miR-3154^34^, miR-548j^35^, and miR-152^36^ in non-small cell lung cancer, abdominal aortic aneurysm, breast cancer, and non-small cell lung cancer, respectively. Interestingly Liu *et al*. noted that together, Tns1 and miR-522-3p reduced NSCLC invasion^33^, where Duan et.al. showed that Tns1 improved NSCLC growth by activation of the Akt/mTOR/RhoA pathway^36^. Another study demonstrated that Tns1 and Tns2 enhances RhoA activity by binding to DIC1 through their domains and showed that a knockout of Tns1 reduced the ability of human umbilical vascular endothelial cells (HUVECs) to participate in cell proliferation, migration, and tube formation as well as decreasing sprouting from aortic rings likely through the Tns1/DIC1/RhoA pathway^37^. To date, our study is the first to show that Tns2 is a target of a microRNA, specifically miR-150-5p.

Angiogenesis is a complex process that not only depends on an appropriate response to angiogenic growth factors and cytokines, but also to the cellular microenvironment, including the extracellular matrix^38^. Such interactions between the cell and its environment are mediated by focal and fibrillar adhesions, comprised of many proteins including integrins, focal adhesion kinase (FAK), and vinculin. These proteins create a dynamic complex that allow the cell to participate in mechanotransduction, encouraging cell adhesion and migration^39^. Focal adhesions are crucial in endothelial cells, promoting endothelial cell elongation and cell-cell junction linearization. When depleted, focal adhesions impair vascular network remodeling and maturation^40^. Here we have shown that alcohol exposure decreases Tns1 and Tns2 expression in BMVECs and leads to a reduction in cell elongation and overall cell sprouting. When overexpressed, Tns1 and Tns2 increase BMVEC sprouting and elongation following alcohol exposure. This effect may, in part, involve interactions between the SH2 domains of Tns1 and Tns2 with FAK^18^. In cancer studies, FAK promotes angiogenesis, cell migration, invasion, and proliferation via cell signaling^41^. Although these findings come from cancer studies, they may inform alcohol-related vascular research. Thus, FAK and its association with Tns1 and Tns2 is a plausible pathway to consider in alcohol-mediated vascular dysfunction. Future studies could test whether alcohol exposure alters FAK activity or its association with Tns1 and Tns2 in BMVECs, and whether disrupting these interactions affects vascular functions such as angiogenesis. Additional miRNA inhibitors predicted to target Tns1 and Tns2 could also be tested in these functional studies. A limitation of this study is that Tns1 and Tns2 overexpression was evaluated *in vitro*; future studies will validate these findings in vivo or in a more physiologically relevant model.

## 5 Conclusion

Our findings show that alcohol exposure impairs BMVEC angiogenic sprouting, which may contribute to cerebrovascular dysfunction following PAE deficits. Increasing Tns1 and Tns2 expression rescued the impaired angiogenic sprouting suggesting that these proteins support developmental angiogenesis. Additionally, we further characterized the role of miR-150-5p as a negative regulator of angiogenesis through its targets Tns1 and Tns2.

