## Supplementary Figures for "Prenatal Alcohol Exposure Downregulates Focal Adhesion Molecules Tensin 1 and Tensin 2 Contributing to Cerebrovascular Dysfunction"

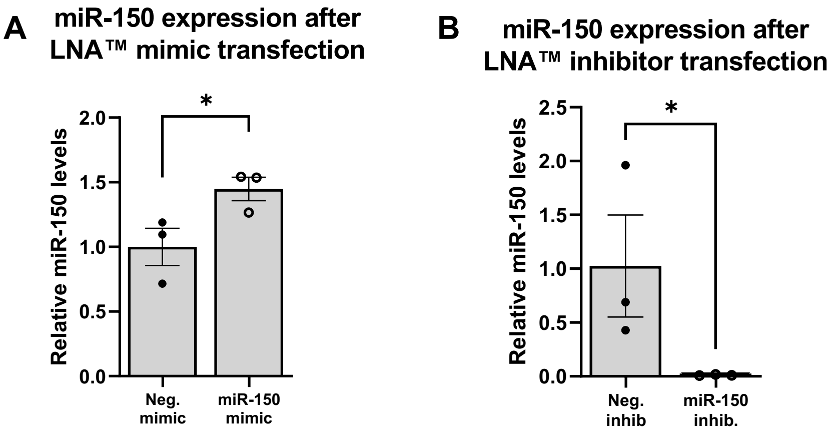


**Supplementary Figure 1:** miR-150-5p levels are significantly altered following overexpression or inhibition in BMVECs. **(A)** miR-150-5p expression in cells following transfection with LNA™ miR-150-5p mimics vs negative controls. RT-qPCR. n=3. p=0.0296. **(B)** miR-150-5p expression in cells following transfection with LNA™ miR-150-5p inhibitors vs negative controls. RT-qPCR. n=3. Error bars=SEM. p=0.0495.

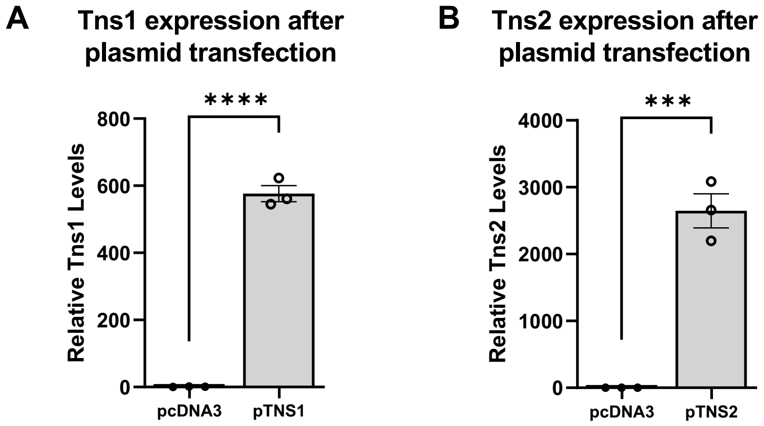


**Supplementary Figure 2:** Tns1 and Tns2 levels are increased following their overexpression in BMVECs. **(A)** Tns1 expression in cells following transfection with a hTns1-expressing plasmid vs pcDNA3. RT-qPCR. n=3. p<0.0001. **(B)** Tns2 expression in cells following transfection with a hTns2-expressing plasmid vs pcDNA3. RT-qPCR. n=3. Error bars=SEM. p=0.0002.
